# Impression cytology is a non-invasive and effective method for ocular cell obtention from babies with Congenital Zika Syndrome: perspectives in OMIC studies

**DOI:** 10.1101/577668

**Authors:** Raquel Hora Barbosa, Maria Luiza B. dos Santos, Thiago P. Silva, Liva Rosa-Fernandes, Ana Pinto, Pricila S. Spínola, Cibele R. Bonvicino, Priscila V. Fernandes, Evandro Lucena, Giuseppe Palmisano, Rossana C. N. Melo, Claudete Cardoso, Bernardo Lemos

## Abstract

**IMPORTANCE:** Noninvasive techniques for obtaining ocular surface cells (neuroepithelial) from babies with Congenital Zika Syndrome CZS - resulting from infection by zika virus (ZIKV) during gestational period (malformations include ocular abnormalities and microcephaly) - remain to be determined.

**OBJECTIVES:** The aim of this study was to describe an optimized impression cytology method for the isolation of viable cells from babies with CZS in satisfactory amounts and quality to enable the application in the context of genome approaches well as morphological and molecular evaluations.

**DESIGN, SETTINGS AND PARTICIPANTS:** In this observational study, ocular surface samples were obtained with a hydrophilic nitrocellulose membrane (through optimized impression cytology method) from twelve babies referred to the Pediatric Service of the Antonio Pedro Hospital, Universidade Federal Fluminense (UFF), Niteroi, Rio de Janeiro, Brazil. Samples were collected with an authorized informed consent from both eyes of eight ZIKV infected babies according to the CZS diagnostic criteria (4 babies with positive PCR for Zika virus in gestation and presence of clinical signs which included ocular abnormalities and microcephaly and 4 babies with positive PCR for Zika virus during gestation but no clinical signs identified) and four unaffected babies (control samples / negative PCR, without clinical signs). Cells were used for microscopy analyses, transcriptomic and proteomic experiments and molecular procedure.

**MAIN OUTCOMES AND MEASURES:** The microscopic features of the conjunctival epithelial cells were described by both direct analysis of the membrane-attached cells and analysis of cytospinned captured cells using several staining procedures, including viability evaluation. In parallel, molecular approaches were performed.

**RESULTS:** On impression cytology, a considerable amount of viable cells were captured. Epithelial basal, polyhedral and goblet cells were clearly identified in all groups. All cases of ZIKV infected babies showed clear morphological alterations (cell keratinization, piknosis, karyolysis, anucleation and vacuolization). Genomic DNA and RNA were successfully isolated from all samples and allowed the establishment of transcriptomic and proteomic studies. Transcriptome analysis showed 8582 transcripts quantified in all samples and 63 differentially expressed genes in ocular cells from the exposed babies. Proteomics analysis allowed the identification of 2080, 2085 and 2086 high confident and unique proteins with at least one unique peptide in the unaffected, exposed to ZIKV and asymptomatic and CZS babies, respectively, being 2062 in common. Multivariate supervised analysis using the total quantitative protein features revealed a clear discrimination between the groups.

**CONCLUSIONS AND RELEVANCE:** Our method proved to be a suitable, fast, and non-invasive tool for detailed and precise morphological analyses with a perspective of application in OMIC studies for clinical and research studies of CZS.

**Key points:** *Question:* Are the ocular surface cells of babies with Congenital Zika Syndrome viable to investigate the association between Zika virus infection during embryogenesis and ocular impairment?

*Findings:* To this date, this is the first study using an approach with perspectives in morphological, molecular and “*OMICs*” research from ocular samples captured by impression cytology of ZIKV infected babies during embryogenesis. The microscopic features of the conjunctival epithelial cells from all ZIKV infected babies showed clear morphological alterations.

*Meaning:* Ocular cell surface capture offers a powerful model for studying the pathways involved in ocular diseases associated with ZIKV.

## Introduction

Zika virus (ZIKV) is an arbovirus of the Flavivirus genus first identified in Uganda - Zika forest, in 1947^1^. In 2015, after an outbreak of acute exanthematic disease, ZIKV was detected in Northeast Region of Brazil^2^,^3^. However, recent studies indicate that the virus circulation in Brazil occurred prior to this epidemic period^4, 5, 6, 7^.

Congenital Zika Syndrome (CZS) was identified due to the increased incidence of congenital defects associated with ZIKV infection. This led to clinical, epidemiological and experimental studies seeking to address the association between congenital defects and ZIKV infection. Furthermore, the World Health Organization (WHO) recognized ZIKV and associated neurological complications as a long-term public health challenge. A global strategic response plan has been issued to enable detection, prevention, care and support in affected areas^8^. Studies were also launched to advance the development of intervention, control and prevention strategies^9^.

Studies on CZS have predominantly involved analysis of brain regions^10^, ^11, 12, 13^ but studies of the ocular system have also been conducted. The studies documented characteristic ocular lesions, such as pigment mottling, macular atrophy, chorioretinal atrophy, horizontal nystagmus and optic nerve hypoplasia and atrophy in the context of ZIKV infection ^14, 15, 16, 17, 18, 19^. Retinal changes occur in about 30 to 40% of cases and anomalies of the development of the eye may occur in several embryogenesis stages such coloboma and ocular structure, including eyelid, cornea, iris, zonula and ciliary body, choroid, retina and optic nerve^20, 21^. Consequently, screening and long-term monitoring of ocular health are crucial to all children with possible congenital ZIKV infection ^22,23^.

Molecular methodologies were established to investigate the association between ZIKV and neurological impairment using induced pluripotent stem cells and embryonic stem cell lines differentiated in neuroprogenitors, neurons, glial cells and into brain organoid structures^24^. However, the use of non-invasive strategies to study ocular cells in ZIKV-infected babies remain to be determined.

Impression cytology of ocular cells is a noninvasive method for external evaluation of ocular lesions^25^. This technique has been developed since the discovery that cells from the eye outside of the epithelial layer could be removed by filter membrane application to evaluate various conditions of ocular surface impairment^26^. This method has been applied to anatomically locate the conjunctiva, quantify goblet cell density, stage squamous metaplasia staging, differentiate bacterial, viral, allergic, degenerative or tumor affections^27,28,29,30, 31,32^.

The aim of this study was to develop ocular impression cytology for the isolation of viable cells in satisfactory amounts and quality to enable the application in the context of genome approaches. OMIC technologies are high-throughput methodologies that have not been coupled with ocular analysis in CZS until this moment. These technologies encompass genomics, transcriptomics, epigenomics, proteomics, and metabolomics, and can provide a global representation of the processes within cells at several levels, contributing to advances in biology and medicine^33^. Here we propose an optimized and noninvasive alternative for obtaining ocular cells from babies with ocular anomalies caused by ZIKV infection during embryogenesis, and its coupling with OMIC applications. Considering that conjunctival cells arise from neurogenic ectoderm^34^ and the well-documented ZIKV neurotropism with affinity for neural progenitor cells ^35, 36, 37, 10, 38, 35^, the methodology presented here may be used to obtain ocular/neuroepithelial cells as potential models of neural cells for CZS research.

## Material and Methods

### Study design and ethic aspects

Twelve babies referred to the Pediatric Service of the Antonio Pedro Hospital, Universidade Federal Fluminense (UFF) were included in this study. All children have been followed by periodical ophthalmological examinations; samples were obtained with an authorized informed consent. This study was approved by Universidade Federal Fluminense Ethics Committee and followed the tenets and guidelines of the Declaration of Helsinki.

Ocular surface samples were collected from both eyes of eight babies according to the CZS diagnostic criteria (Group A: Patients 1 to 4 = positive PCR for Zika virus in gestation and presence of clinical signs which included ocular abnormalities and microcephaly (ZIKV/CZS); Group B: Patients 5 to 8 = positive PCR for Zika virus during gestation but no clinical signs identified; ZIKV) and four unaffected babies (Group C; control samples: Patients 9 to 12 = negative PCR, without clinical signs; CTRL).

### Impression cytology and ocular surface cells capture

A local anesthetic (Proxymetacaine hydrochloride, 0.5% w/v, eye drops, solution.) was instilled into the eye before obtaining the ocular surface samples. The samples were collected with a sterilized 0.45µm, 47mm white plain hydrophilic nitrocellulose membrane (Millipore Sigma®, catalogue number HAWP047S0). Each circular membrane was cut into four strips measuring 0.75cm wide and 4.5 long approximately (Figure 1).

The method here described does not use tweezers or pediatric lid speculum for sample collection. A stem of the membrane is used as a collection support. The strip ends were rounded and bent at approximately 1cm to facilitate printing on the ocular surface and comprises the capture area of ocular cells (Figure 1). The strip end was then pressed on to the inferior bulbar conjunctiva for approximately five seconds. The strip stem was discarded after the collection by using a sterilized scissors (Figure 1).

**Figure 1.**
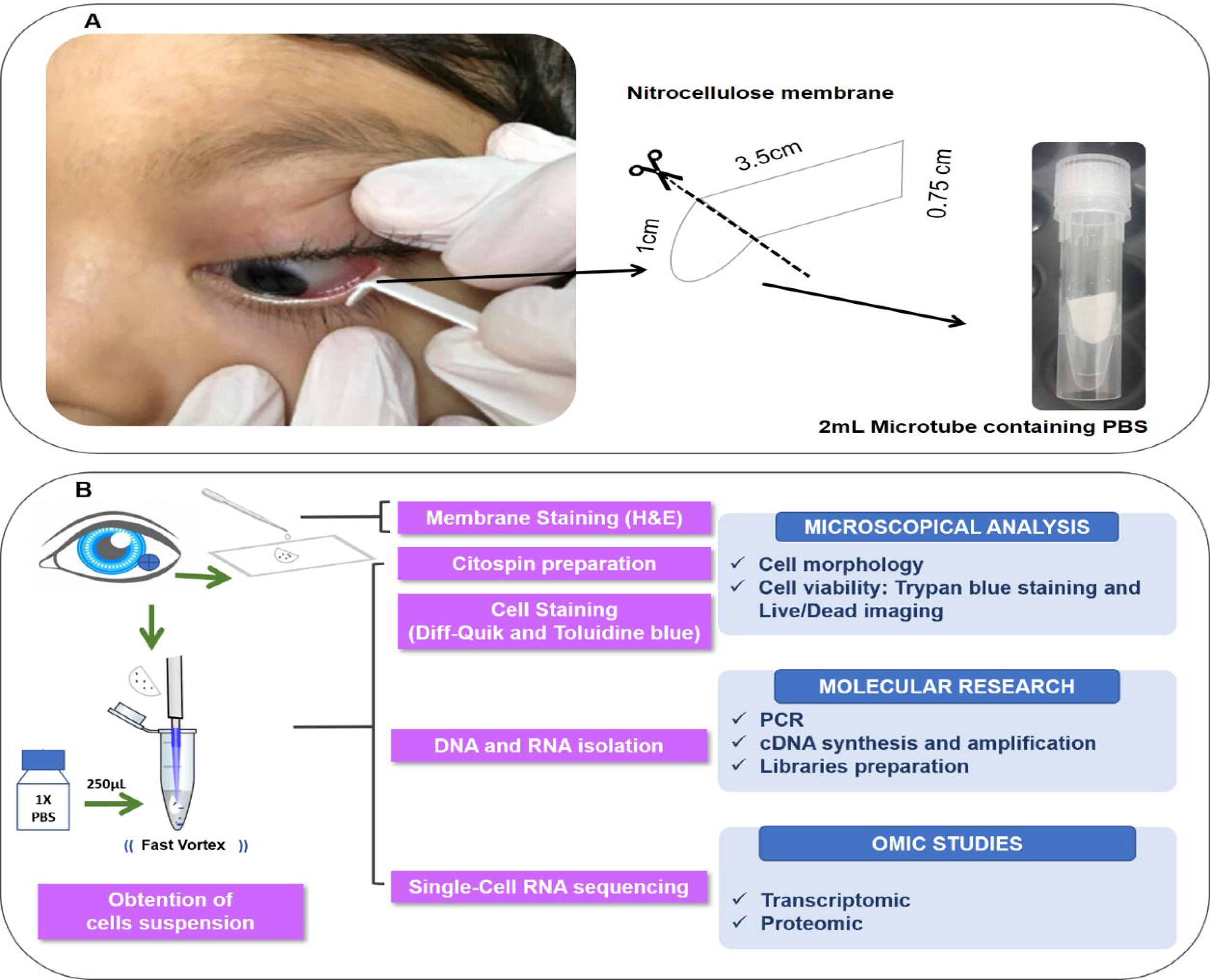
Overview of the non-invasive strategy (impression cytology method) used to collect ocular cells from children with Congenital Zika Syndrome and applied methodologies. **A.** Impression cytology was optimized (without use of tweezers and pediatric lid speculum) for sample collection. The membrane model has a rounded apex and a long support base that provides a correct positioning and safer method for the collection of baby ocular samples. **B.** Methodologies applied to different studies of captured cells.

### Cell collection and storage

The cell capture area of the filter membrane was immediately placed in a 1.5 mL tube on ice containing 250 µL 1X PBS (Phosphate Buffered Saline, pH7.4 - TermoFisher Scientific, catalog number: 10010023). Tubes containing filter membranes were rapidly vortexed to allow release of adhered cells (Figure 1). Cell suspensions were then used for microscopy analyses or aliquoted in 0.2 mL tubes and stored for additional experiments. Specifically, we aliquoted 9 μL for Transcriptomic experiments - Single Cell RNAseq (stored at - 80°C); 100 μL for Proteomic experiments (stored at - 80°C) and 100 μL for DNA isolation (stored at 4°C).

### Cell analysis and Microscope Image Acquisition

#### Cell Viability

Viability of the collected cells was determined by the trypan blue exclusion test [9μL of the suspension cells plus 1μL of 0.4% trypan blue solution (TermoFisher Scientific, catalog number: 15250061)]. Viable cells were counted in a Neubauer chamber. Images were acquired using on Axio microscope with 20x/0.35 and 40x/0.55 Zeiss A-Plan objectives (Carl Zeiss, Jena, Germany, http://www.zeiss.com) and Q-Capture PRO 7 software (Surrey, BC, Canada, www.qimaging.com).

Additionally, cytospin procedure was used for allows the concentration of single cells in suspension on a microscope slide. Cytocentrifuged preparations (200 μL of cell suspension/slide) were obtained in a Cytospin 4 Shandon (Thermo Scientific Corporation, Waltham, MA) at 800 rpm for 5 minutes at room temperature and stained with LIVE/DEAD^TM^ viability/cytotoxicity kit (ThermoFisher Scientific, catalog number L3224) as the manufacturer’s instructions. This kit contains a mixture of fluorescent stains (calcein-AM and ethidium homodimer-1) which discriminates live from dead cells by simultaneously staining with green-fluorescent calcein-AM to indicate intracellular esterase activity and red-fluorescent ethidium homodimer-1 to indicate loss of plasma membrane integrity. Analyses were performed on a fluorescence microscope (BX-60, Olympus, Melville, NY, USA) using U-MWB FITC/Texas red filter (488–570 nm excitation wavelengths), which allows simultaneous visualization of both markers.

#### Cell Morphology

To evaluate microscopic features of the captured cells such as morphological types and possible alterations we then used another collection membrane directly stained with hematoxylin and eosin after fixation for 10 min in a fixation solution (100 mL of 70% ethanol, 5 mL of glacial acetic acid and 5 mL of 37% formaldehyde solution – all solutions are from Merck Millipore). Samples were hydrated with distilled water for 5 minutes, immersed in Harris' hematoxylin for 2 minutes, washed with tap water for 15 minutes, counterstained with eosin for 30 seconds, washed with tap water for 5 minutes, and then dehydrated in 70%, 80%, 95%, and 100% ethyl alcohol (rapid immersion for 5 seconds each). After these steps, samples were immersed in xylene (ten successive immersions for 5 minutes each) and mounted on slides cover-slipped using Entellan mounting medium (Merck, Millipore). Images were acquired using on Axio microscope with 20x/0.35 and 40x/0.55 Zeiss A-Plan objectives (Carl Zeiss, Jena, Germany, http://www.zeiss.com) and Q-Capture PRO 7 software (Surrey, BC, Canada, www.qimaging.com).

Another cytocentrifuged preparation was also stained for morphological analyses. For this, cell suspensions obtained as above were fixed in 4% paraformaldehyde and cytocentrifuged (Cytospin 4 Shandon, Thermo Scientific, 1200 rpm, 10 minutes). Slides of captured cells (n=3 patients for each group) were prepared in quadruplicate. For each pair of slides, one was stained with a Diff-Quik kit, as the standard procedure, and the other one with 0.5% toluidine blue O solution (Fisher Scientific) for 5 minutes. Cells were analyzed on a BX-60 Olympus microscope,

#### Molecular analyses

To evaluate the integrity and quality of the genomic material of captured ocular surface cells, DNA was isolated according to QIAamp DNA mini kit (Qiagen, Valencia, CA), which is indicated for swabs, body fluids or washed cells. Thereafter, we used a pair of primers to amplify a region (exon 4) of the *MCPE2* gene that possess a significant role in embryonic development. The forward: U-GGA AAG GAC TGA AGA CCT GTA AG and Reverse R-CTC CCT CCC CTC GGT GTT TG (fragment size = 372pb) primers were used and PCR conditions with modifications were applied according to previous report^39^. The PCR was performed in a total volume of 25µL, including 1µL DNA template, 1µL of each primer (10 μM), 13.8µL distilled ddH2O, 5.0µL buffer, 2.5 µL MgCl_2_, 0.5 µL dideoxynucleotide and 0.2µL of the Platinum Taq DNA polymerase (Platinum^TM^ Taq DNA polymerase, Catalog number: 10966026; Thermo Fisher Scientific). Touchdown program was used in the Veriti 96 Applied Biosystems PCR thermal cycler and all samples were denatured at 95°C for 2min and 30sec, followed by: 25 cycles of 94°C for 30sec, 65°C for 30sec, and 72°C for 1min45sec, followed by 10 cycles of 94°C for 30sec, 51°C for 30sec and and 72°C for 1min30sec then a final extension 72°C for 5min. For each sample, 5μL of the final PCR product was checked by 1% agarose gel electrophoresis.

For transcriptomic experiments 9μL of suspension solution (after fast vortex of capture membrane – Figure 1) containing a maximum number of 150 cells were processed through the Single Cell RNASeq Service (SingulOmics^40^/ Novogene^41^) which included cDNA synthesis and amplification, library preparation, sequencing (10 million paired-end reads) and data analysis.

Here, total RNA quality assessment in the samples including preparation of the Single Cell RNA library was performed by SingulOmics^40^ using the 2100 Bioanalyzer^42^ - a microfluidic platform for the electrophoretic separation of biomolecules - extremely useful for identifying contaminated and degraded RNA (Figure 4).

**Figure 4.**
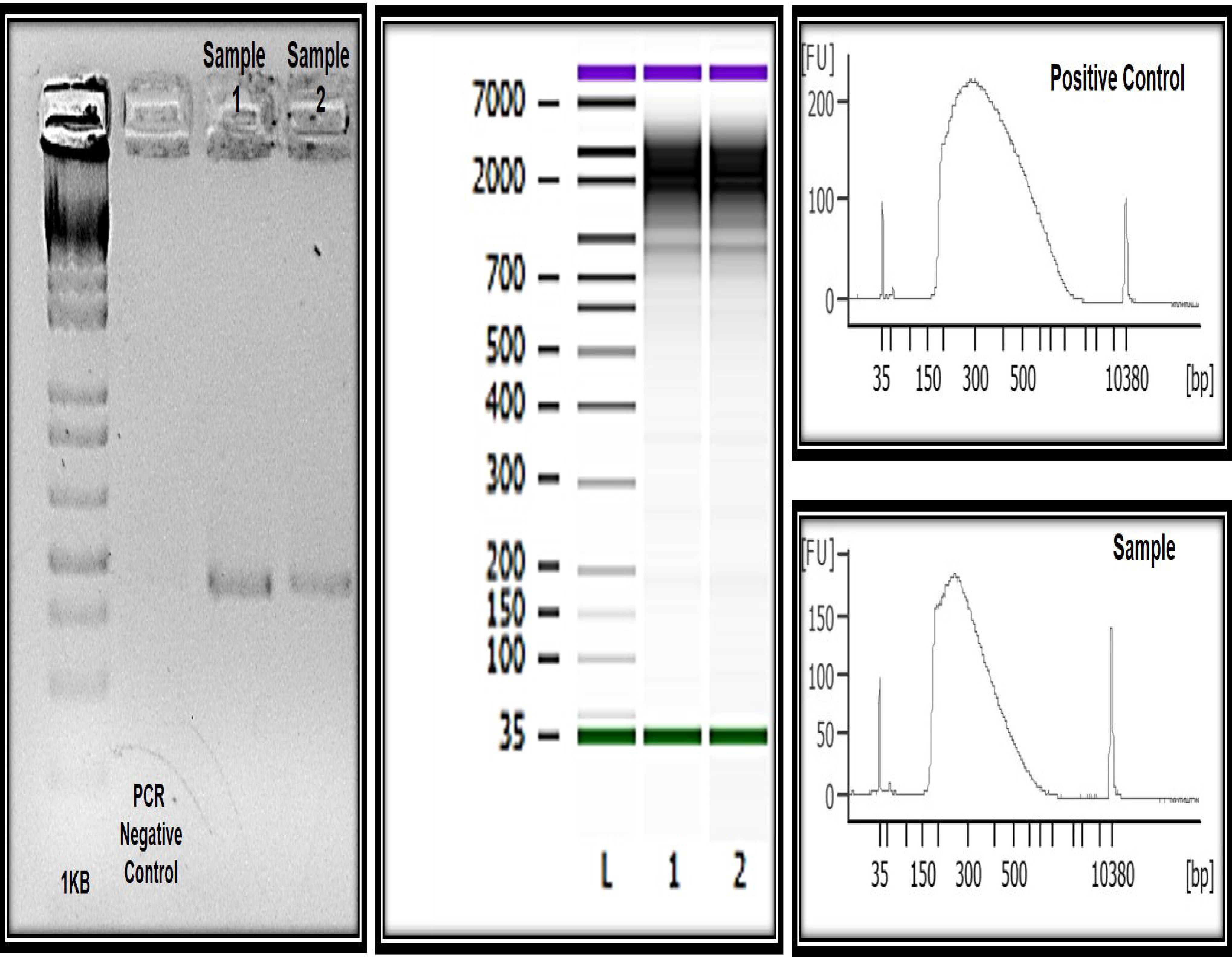
Molecular and “OMIC” research perspectives. **A.** Electrophoresis in 1.5% agarose gel and the amplification of a 350bp fragment correspondent to amplicon 4 of the *MCPE2* gene in DNA of the ocular surface cells. **B^*^.** Quality and integrity RNA analysis. L= ladder, 1 and 2= CZS children. **C\***. Quality control libraries - comparative concentrations for positive control (of the used method) and healthy child (control sample of this study). **\***Provided by Singulomics^©^.

For proteomics experiments proteins were extracted from the membrane, digested with trypsin and analysed in a data-dependent manner by nanoflow liquid chromatography coupled to high accuracy and resolution mass spectrometry, Orbitrap Fusion tribrid, Thermo Fisher. Raw data were searched using Sequest database search engine using reviewed Uniprot human protein database. Protein identifications were filtered with less than 1% FDR.

## Results

### Clinical aspects

Samples were collected from both eyes of eight boys and four girls’ babies with 21 months median age (Table 1). ZIKV infected babies according to the CZS diagnostic criteria (4 babies with positive PCR for Zika virus in gestation and presence of clinical signs which included ocular abnormalities and microcephaly – ZIKV infection predominantly in the first trimester), 4 babies with positive PCR for Zika virus during gestation (occurring in the second and third trimester) but no clinical signs identified and 4 unaffected babies (control samples / negative PCR, without clinical signs).

**Table 1.** Clinical data.

| Subject's ID | Gender | Status (ZIKV exposition during pregnancy) | Status (for microcephaly) | Vision impairment | Age at samples collection (in months) | ZIKV infection symptoms and PCR |
| --- | --- | --- | --- | --- | --- | --- |
| #1 | M | Exposed | Affected | Yes | 21 | 1<sup>o</sup> trimester |
| #2 | M | Exposed | Affected | Yes | 22 | 1<sup>o</sup> trimester |
| #3 | M | Exposed | Affected | Yes | 19 | 2<sup>o</sup> trimester |
| #4 | F | Exposed | Affected | Yes | 27 | 1<sup>o</sup> trimester |
| #5 | M | Exposed | Non-affected | No | 21 | 2<sup>o</sup> trimester |
| #6 | M | Exposed | Non-affected | No | 20 | 2<sup>o</sup> trimester |
| #7 | F | Exposed | Non-affected | No | 24 | 3<sup>o</sup> trimester |
| #8 | F | Exposed | Non-affected | No | 24 | 3<sup>o</sup> trimester |
| #9 | M | Non-exposed | Non-affected | No | 9 | - |
| #10 | F | Non-exposed | Non-affected | No | 21 | - |
| #11 | M | Non-exposed | Non-affected | No | 24 | - |
| #12 | M | Non-exposed | Non-affected | No | 19 | - |

### Cell visualization, counting, distribution and morphological aspects

Here we developed a membrane model for cell collection with a rounded apex and a long base of support that provided a correct positioning for the capture time and for fixing and staining. We used the membrane extremity as collection support and then discarded not requiring the use of tweezers and pediatric lid speculum (Figure 1).

The impression cytology with nitrocellulose membrane model we developed here is effective for ocular surface cells capture. We observed the presence of cells in all collection membranes. In only 9µL of cell suspension after fast vortex (from an initial total of 250 µL), the number of cells retrieved ranged from 15 to ~150 cells. Most cells remained attached to the membrane.

Cell viability evaluations by trypan blue test of captured cell suspensions showed that most cells (> 95%) were viable immediately after collection in all groups and different epithelial and goblet cells were observed in Neubauer *chamber*. In addition, live cells were imaged by intense, uniform green fluorescence (Figure 2A) while dead cells fluoresced orange-red after staining with a live/dead viability kit (Supplementary Figure 1). We detected the presence of viable cells adhered to the collection membrane (attached to the nitrocellulose fibers) 5h after application of impression cytology (Figure 2).

**Figure 2.**
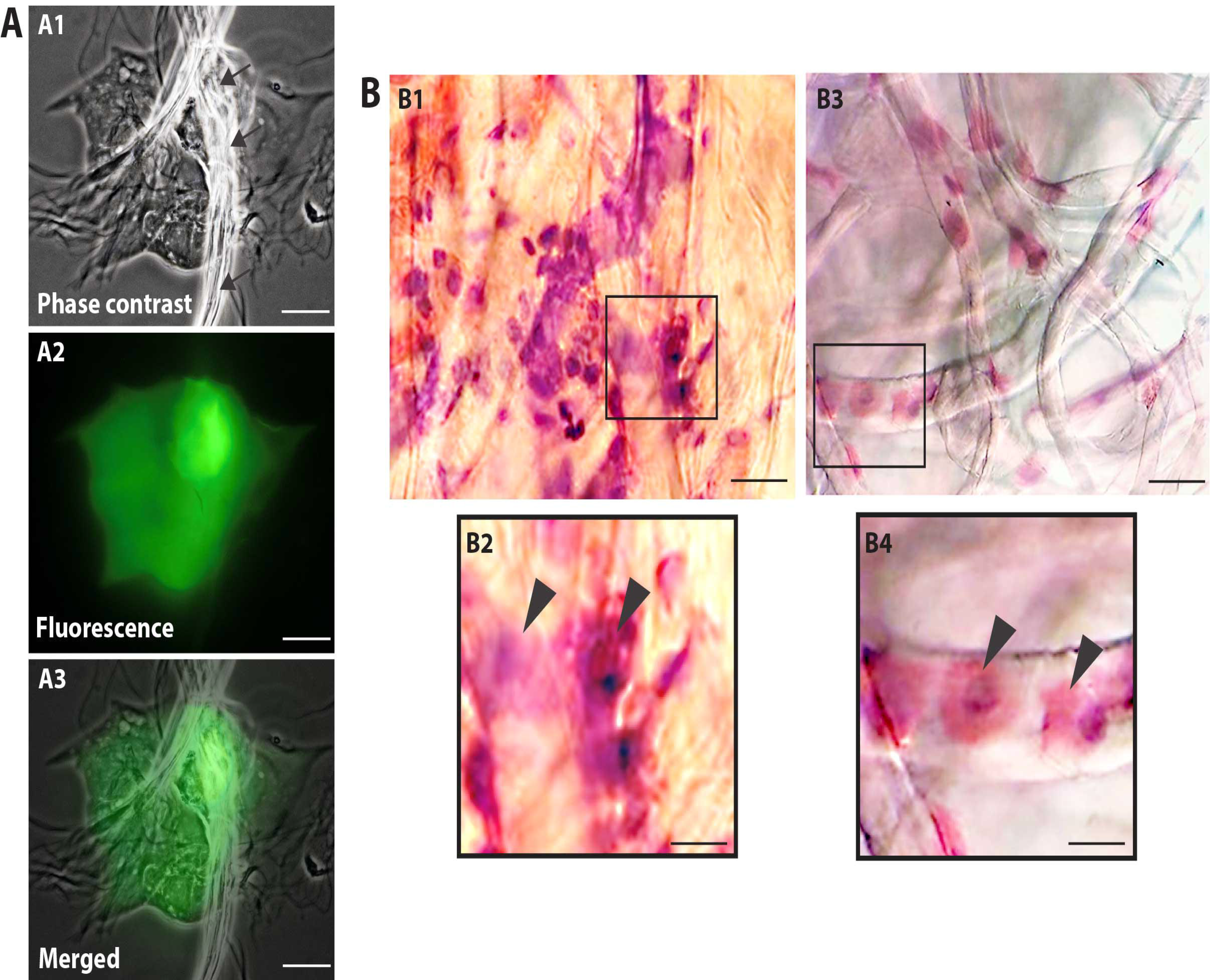
Nitrocellulose membrane fibers impregnated by ocular surface cells from CZS patient (A, B1, B2) and uninfected controls (B3 and B4). **A.** Representative live cells are shown by phase contrast (A1) and fluorescence microscopy (A2). An overlay of these two images is seen in (A3). Viable cells fluoresce in bright green after staining with LIVE/DEAD^®^ cell viability/cytotoxicity assay. Cells were imaged 5h after collection**. B.** Representative membranes directly stained with hematoxylin/eosin. Membrane-attached cells are indicated by arrowheads in higher magnification in (B2) and (B3). Scale bars, 20 μm (A1-A3); 70 μm (B1); 35 μm (B2); 50 μm (B3); 25 μm (B4).

Microscopic analysis of the stained membranes (Figure 2B), showed presence of varying amounts of cells in all of them. Individuals infected with ZIKV (Figure 2B1 and B2) showed apparently more cells attached to the membrane when compared with control subjects (Figure 2B3 and B4). Some impression areas stained more intensely likely due to multilayering of the cells. The morphologic evaluation was impaired in these regions. Of note, we observed that the eyes with excessive tearing had worse results in the capture of cells; however, this did not affect both the collection and additional procedures.

Morphological cell analyses were performed after citospinning the cell suspensions which facilitated adhesion of the cells on the slides and resulted in enhanced visualization of cell features (Figure 3). The following cell types, characteristic of the conjunctiva, were identified in all groups: epithelial basal cells, epithelial polyhedral cells and goblet cells (Figure 3A). Epithelial cells of the basal layer were seen individually or in small clusters, with a round to oval shape and a central nucleus and scant cytoplasm (Figure 3A2 and A3). Basal cells stained more strongly compared to other epithelial cells. Intermediate and more superficial epithelial cells were recognized by their polyhedral and abundant cytoplasm with a small and central nucleus (Figure 3A4 and A5). Goblet cells were identified by their typical morphology – an eccentric nucleus and a pale cytoplasm in their apical region (Figure 3A6 and A7). When the groups were compared, morphological changes were clearly detected in cells collected from ZIKV infected patients, predominantly in those with clinical signs (CZS), compared to uninfected controls. These included nuclear and cytoplasmic alterations such as mild to moderate keratinization (Figure 3B1-B4), piknosis (Figure 3B4 and B5), karyolysis (Figure 3B2 and B3), anucleation (Figure 3B6) and cytoplasmic vacuolization (Figure 3B7 and B8).

**Figure 3.**
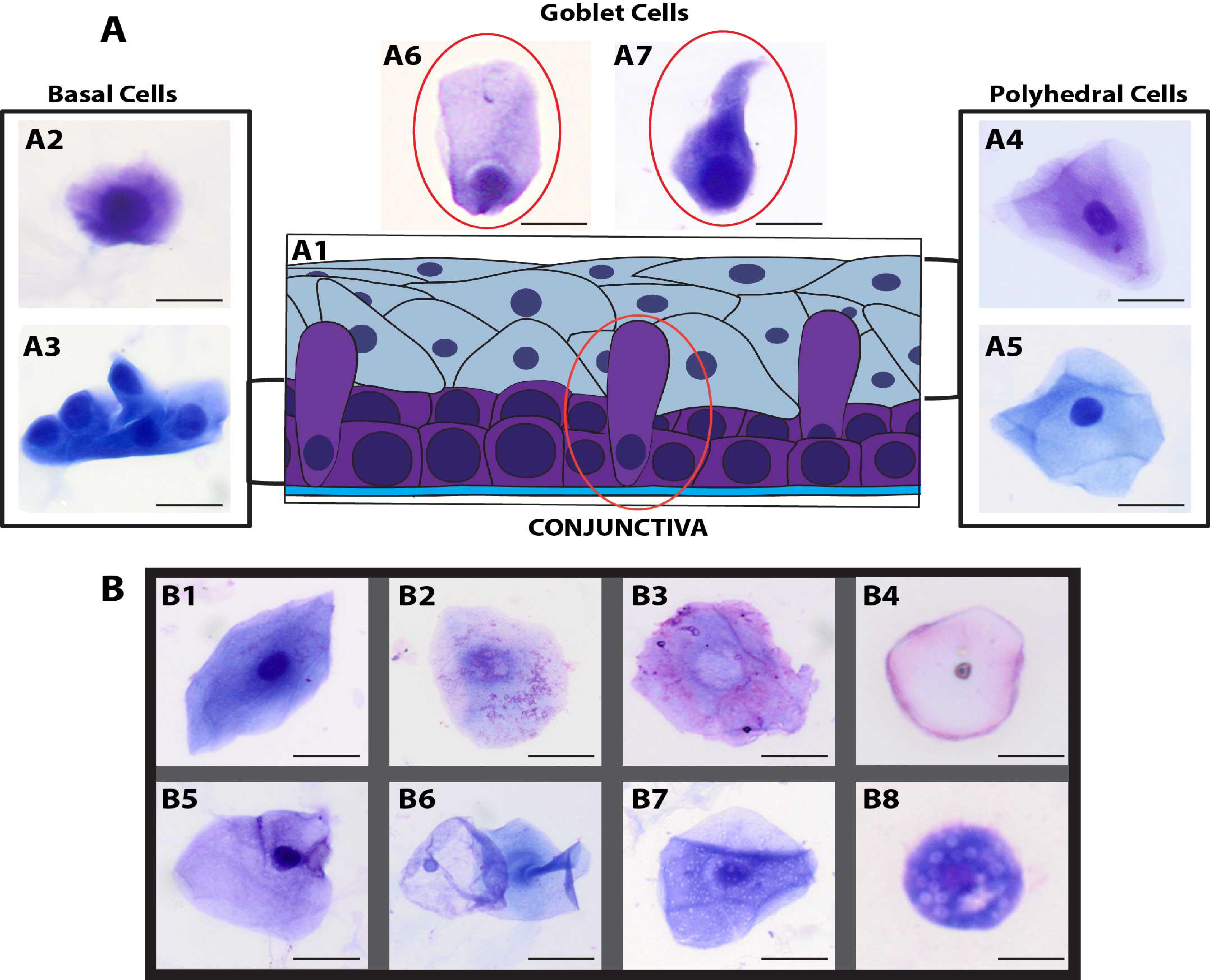
Morphological analyses **A.** Different ocular surface cells collected from impression cytology. **A1**. Human conjunctiva cell types pattern. Representative basal, polyhedral and goblets cells from uninfected (A2-A4, A6) and ZIKV-infected children with no clinical signs (A5, A7) show normal morphology. **B.** Morphological changes observed in cells from ZIKV infected children without clinical signs (B1, B2) and CZS children (B3-B8). Cell changes included mild to moderate keratinization (B1-B4), piknosis (B4 and B5), karyolysis (B2 and B3), anucleation (B6) and vacuolization (Figure B7 and B8). Cytocentrifuged preparations were stained with Diff-Quik (A2, A4, A6, B1-B4) or toluidine blue (A3, A5, A7, B5-B8). Scale bars, 5 μm (A6, A7); 10 μm (A2, B2-B5); 20 μm (A3-A5, B1, B7); 25 μm (B6).

### Molecular applications and genomic analyses

Genomic DNA was successfully isolated from all samples. The amount of genomic DNA ranged from 10ng/μL to as much as 70ng/μL, with good integrity, and sufficient for successful PCR reactions. A unique fragment of 372bp correspondent to partial region of exon 4 of the *MCPE2* gene was detected in all samples (Figure 4). We also obtained whole and viable cells in good quality for downstream applications using transcriptomic, epigenetic, or proteomic approaches (Figure 4 and unpublished data). As an example, RNA preparations were obtained and proceed with RNA-seq. Given the small number of cells in some preps, we conducted whole transcriptome PCR amplification prior to sequencing. Sufficient RNA and transcriptome libraries were obtained for all samples. Transcriptome analysis showed 8582 transcripts quantified in all samples (5 reads or more). Relative to the unaffected group (CTRL) group, we observed 63 differentially expressed genes (pvalue<0.001) when ocular cells from the exposed babies (ZIKV/CZS and ZIKV; Figure 5 and unpublished data).

**Figure 5.**
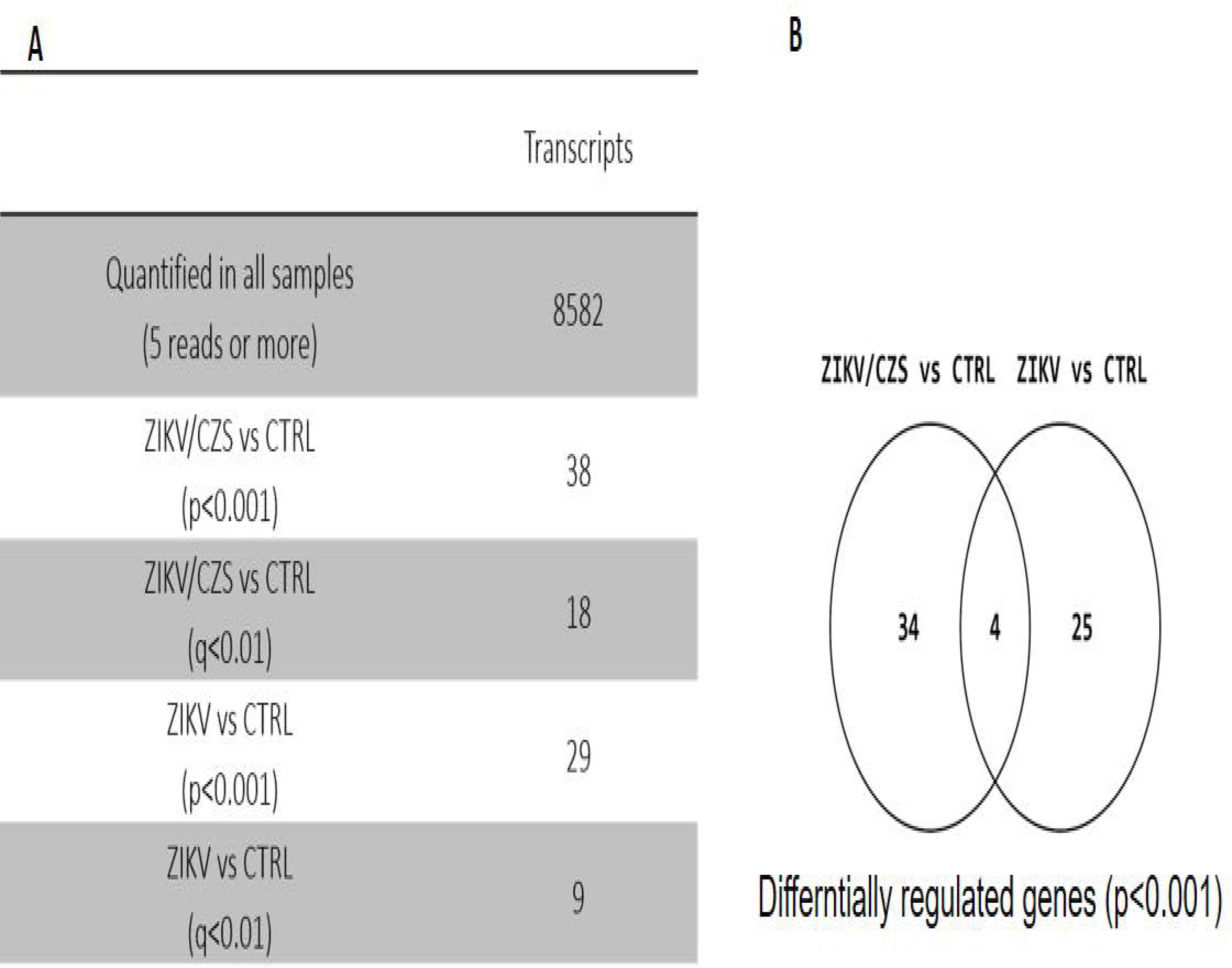
Ocular surface cells gene expression analyses by RNA-Seq. **A.** Descriptive analysis table. **B.** Transcripts quantified and regulated. Venn diagram of DRG (p<0.001).

Large scale quantitative mass spectrometry-based proteomics analysis allowed the identification of 2080, 2085 and 2086 high confident and unique proteins with at least one unique peptide in the CTRL, ZIKV and ZIKV/CZS conditions, respectively, being 2062 in common; Table 2 and unpublished data). Multivariate supervised analysis using the total quantitative protein features revealed a clear discrimination between the CTRL and ZIKV/CZS groups.

**Table 2.** Number of of identified proteins by mass spectrometry-based proteomics analysis.

| **Condition** | **N° of identified proteins** |
| --- | --- |
| **CTRL** | 2080 |
| **ZIKV** | 2085 |
| **ZIKV/CZS** | 2086 |
| **Common between the three conditions** | 2062 |

## Discussion

Impression cytology has been shown to be a simple and reproducible technique that can be successfully performed in preterm or term infants^43^. However, some authors have reported difficulties in obtaining adequate samples of infants^44,45^. The methods previously described mostly used tweezers and/ or pediatric lid speculum for collection of samples. Here we optimize the technique and discard the use of invasive apparatus, enabling a safer and more effective method for the collection of baby ocular samples. The protocols can be straightforwardly performed with sufficient training and easily scaled for analyses of larger clinical populations and under a variety clinical context.

### Ocular surface cells: perspective applications in CZS studies

Approximately 50% of children born with CZS and microcephaly present serious eye diseases^46^. Furthermore, the ZIKV has the potential to survive for long periods in ocular tissue and potentially cause outcomes that will only be manifested later in life^47^. The human ocular surface, a specialized region derived from neurogenic ectoderm which includes the corneal, limbal, and conjunctival stratified epithelia, play an essential role in ocular system^48^. Due to zika virus neurotropism for infect neural cells in human embryonic development, the ocular cells obtained in this study may represent an adequate model for analysis of molecular alterations resulting from ZIKV virus and neuronal cells interaction. Moreover, since viral ZIKV RNA may be present in ocular fluids (in tears and lacrimal glands)^49, 50, 51^, in our study we provide a methodology for cell capture with different perspectives of application. For example, the technique is applicable to the immunolocalization of a wide range of proteins, including detection of ZIKV antigens; to viral analysis through Real Time PCR and ultrastructural microscopy. Studies of the cellular anatomy, physiology and molecular aspects of the ocular surface are essential for understanding ZIKV-associated ocular and neurologic disorders.

Since ZIKV infection has been related with central nervous system abnormalities, the investigation of alterations in genes associated with syndromes microcephaly and other syndromes is crucial. The *MECPE2* gene, for instance, has been linked to Rett syndrome and Angelman syndrome, X-linked mental retardation, neonatal encephalopathy (severe brain dysfunction in males who live only into early childhood), some cases of autism and systemic lupus erythematosus^52^. Here, with neuroepithelial cells, obtained non-invasively, we have successfully standardized a molecular study protocol for the *MCPE2* gene and can be optimized for several other molecular studies involving other genes of interest investigating the ZIKV and microcephaly association.

Molecular and cellular events, fundamental to embryological development, postnatal maturation, and maintenance of the ocular surface, are specifically regulated through advanced gene expression mechanisms. Several studies suggest a significant discrepancy between transcription and protein levels in specific cells, indicating that mechanisms related to regulation of alternative splicing, transcript stability, translation efficiency, protein stability also participate intrinsically in gene expression^53,54^. With the introduction of transcriptomic and proteomics tools we can compare the findings between the corresponding transcript and protein levels. In this study, we showed cells to be viable both for transcriptomic research via RNAseq technology and for proteomic validation. The quality of the RNA and libraries obtained in the study of transcriptome profiles is crucial for generating accurate and informative results. Transcriptome and proteomic profiling revealed differences between exposed and controls babies. However, this work focuses objectively on the detailed report of the methods performed to obtain viable cells from infants exposed and not exposed to zika virus with appropriate conditions for the establishment of molecular and OMICs procedures. The large-scale data generated from the different high throughput analyses (transcriptomics and proteomics) are detailed in complementary and separate works. Thus, the analysis of differentially expressed genes identified and CZS associated are described in the Barbosa *et al*., 2019 manuscript (submitted) and proteomic data are described in the Rosa-Fernades and Barbosa *et al*., 2019 manuscript (submitted).

Despite their importance, many questions about the genetic characteristics of the conjunctival cells, mainly goblet cells are understudied and deserve further exploration^55^. Moreover, the molecular and morphological aspects of human conjunctival stem cells also have not been clearly elucidated^41^.

The strategies here used enabled clear detection of morphological changes in cells from ZIKV-infected patients such as cytoplasmic keratinization and nuclear alterations as observed in other ocular disorders using cytology impression approach^56^. To this date, this is the first study using an approach with perspectives in morphological, molecular and “*OMICs*” research from ocular samples captured by impression cytology of babies with CZS. Studies of mechanisms involved in CZS in ocular cells require rapid, highly reproducible, and accurate quantification and can be successfully achieved with impression cytology. Ocular cell surface capture offers a powerful model for studying the pathways involved in ocular diseases associated with ZIKV.

## Conclusion

The impression cytology with nitrocellulose membrane model developed in this study is safe and effective method for babies ocular surface cells collection and can be applied to morphological, molecular and “OMIC” research in CZS studies.

## Acknowledgements

We thank to the Zika Study Group of the Clinical Research Unit from the Universidade Federal Fluminense especially to the patients and their families and to Mrs. Angela Brum for the local assistance. We thank to Dr. Hector Seuanez and Dr. Miguel Moreira, researches from the INCA - Genetic Program, for the laboratorial facilities, discussion and for helpful suggestions. We also thank to MD. Fernando Agarez for help us with pathological reports and to Mr. Carlos Silva for the logistical support to the study.

## Author contributions

### Concept and design

RHB, EL, CC

### Acquisition, analysis, or interpretation of data

RHB, MLBS, TPS, LRF, AP, PSS, CRB, EL, GP, RCNM, CC, BL.

### Drafting of the manuscript

RHB, RCNM, CC, BL.

### Critical revision of the manuscript for important intellectual content

RHB, RCNM, CC, BL.

---

**Figure S1.**
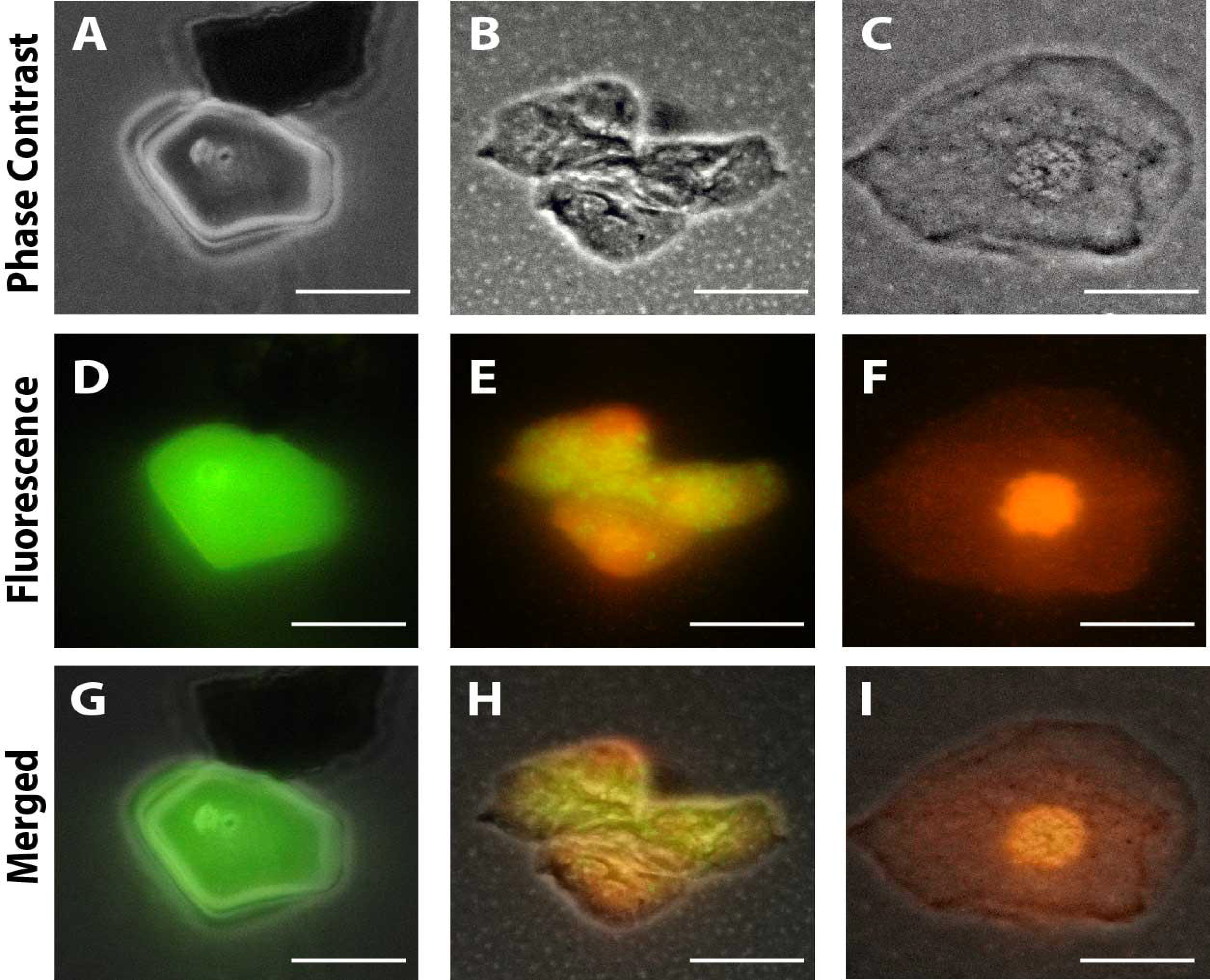
Representative viable and non-viable cells recovered from the conjunctiva of children using an impression cytology method. Live cells fluoresce green whereas dead cells with compromised membranes fluoresce red-orange after staining with LIVE/DEAD^®^ cell viability/cytotoxicity assay. (A, D, G) are from uninfected patients; (B, E, H) are from CZV patients (positive PCR during gestation) with no clinical signs; (C, F, I) are from CZV children (positive PCR) with diagnosed clinical signs (ocular abnormalities and microcephaly). Scale bars, 10 μm (A, D, G); 20 μm (B, H, C); 20 μm (C, F, I). Cells were imaged 5h after collection.

